# Endogenous sterol synthesis is dispensable for *Trypanosoma cruzi* epimastigote growth but not stress tolerance

**DOI:** 10.1101/2022.05.03.490464

**Authors:** Peter C. Dumoulin, Joshua Vollrath, Madalyn Won, Jennifer X. Wang, Barbara A. Burleigh

**Affiliations:** Harvard T.H. Chan School of Public Health, Department of Immunology and Infectious Diseases, Boston, MA, USA; Heidelberg University, Institute for Pharmacy and Molecular Biotechnology, Germany; Harvard University, Harvard Center for Mass Spectrometry, Cambridge, MA, USA

**Keywords:** CYP51, squalene epoxidase, *Trypanosoma cruzi*, ergosterol biosynthesis inhibitors, sterols

## Abstract

In addition to scavenging exogenous cholesterol, the parasitic kinetoplastid *Trypanosoma cruzi* can endogenously synthesize sterols. Similar to fungal species, *T. cruzi* synthesizes ergostane type sterols and is sensitive to a class of azole inhibitors of ergosterol biosynthesis that target the enzyme lanosterol 14α-demethylase (CYP51). In the related kinetoplastid parasite *Leishmania donovani*, CYP51 is essential, yet in *Leishmania major*, the cognate enzyme is dispensable for growth; but not heat resistance. The essentiality of CYP51 and the specific role of ergostane-type sterol products in *T. cruzi* has not been established. To better understand the importance of this pathway, we have disrupted the CYP51 gene in *T. cruzi* epimastigotes (*ΔCYP51*). Disruption of CYP51 leads to accumulation of 14-methylated sterols and a concurrent absence of the final sterol product ergosterol. While *ΔCYP51* epimastigotes have slowed proliferation compared to wild type parasites, the enzyme is not required for growth; however, *ΔCYP51* epimastigotes exhibit sensitivity to elevated temperature, an elevated mitochondrial membrane potential and fail to establish growth as intracellular amastigotes *in vitro*. Further genetic disruption of squalene epoxidase (*ΔSQLE*) results in the absence of all endogenous sterols and sterol auxotrophy, yet failed to rescue tolerance to stress in *ΔCYP51* parasites, suggesting the loss of ergosterol and not accumulation of 14-methylated sterols modulates stress tolerance.

## Introduction

Sterols are integral components of eukaryotic membranes that influence membrane fluidity and architecture (Dufourc, 2008). While some eukaryotic parasites fulfill their sterol requirements solely through cholesterol scavenging (e.g., *Plasmodium* sp.), kinetoplastid protozoan parasites (e.g., *Trypanosoma* sp. and *Leishmania* sp.) maintain an endogenous sterol synthesis pathway that, similar to yeast, produces ergostane-type sterols (Liendo et al., 1999). Consequently, anti-fungal compounds that target endogenously synthesized sterols in the membrane (i.e., polyenes) or inhibit intermediate steps in sterol synthesis (e.g., azoles) have been investigated for their ability to clear certain parasitic infections.

Azoles are a class of ergosterol biosynthesis inhibitors that block the activity of lanosterol 14α-demethylase (CYP51), an enzyme in the sterol synthesis pathway that mediates the demethylation of lanosterol (Figure 1A). A substantial body of pre-clinical data demonstrated the ability of azoles to kill *T. cruzi* parasites (Docampo et al., 1981; McCabe et al., 1984; Buckner and Urbina, 2012), yet, clinical trials (Molina et al., 2014; Torrico et al., 2018) and treatment studies in mouse models of acute and chronic Chagas disease (Francisco et al., 2015; Khare et al., 2015) reveal that azoles suppress *T. cruzi* infection but do not eliminate parasites following treatment. These data suggest that inhibition of CYP51 does not always lead to parasite death; therefore, understanding the contextual importance of endogenous sterols synthesis and the consequences of blocking CYP51 activity will aid in evaluating partner therapies.

**Figure 1:**
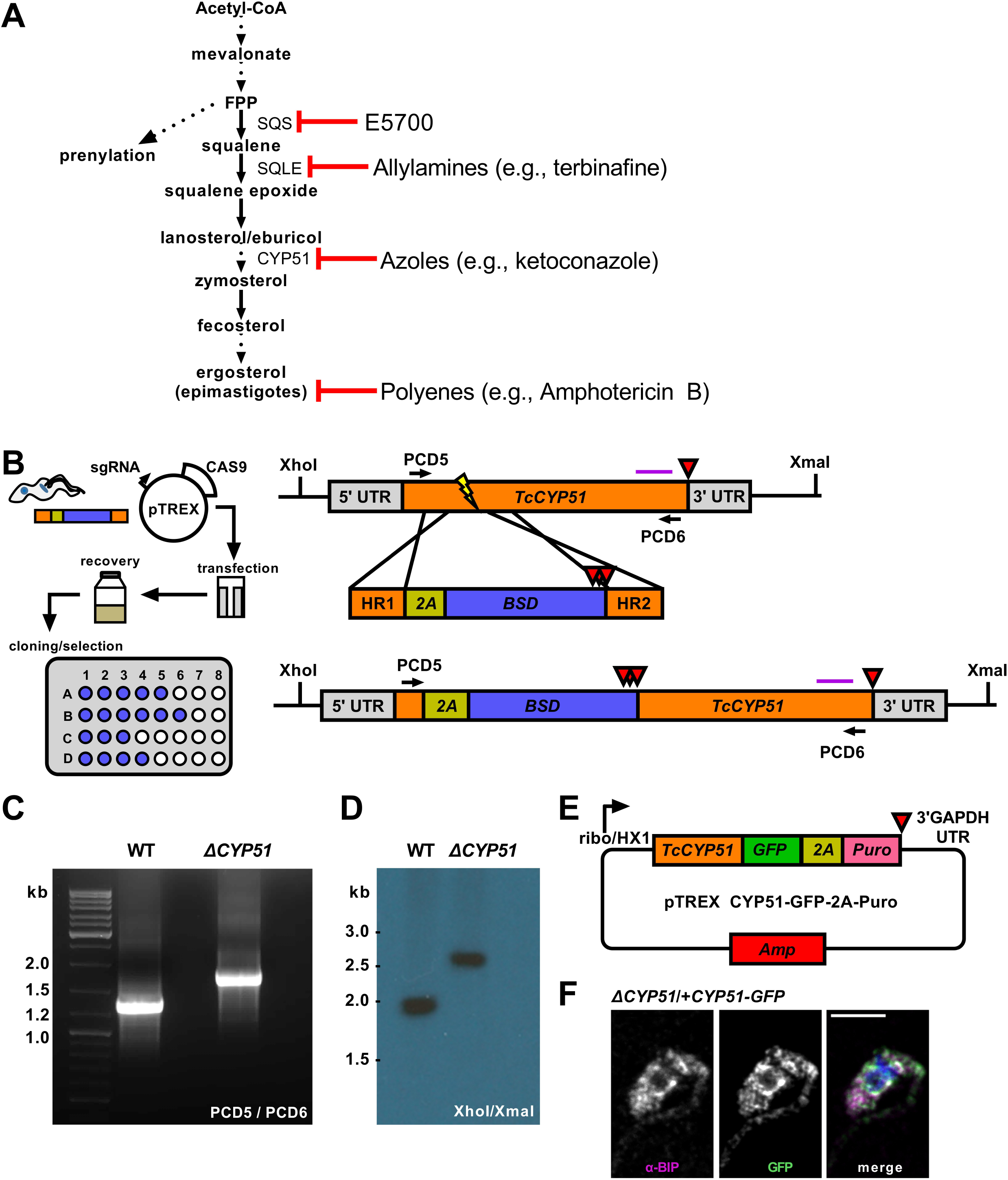
**(A)** Outline of endogenous sterol synthesis in *T. cruzi* epimastigotes. Dashed arrows indicate the omission of steps for clarity. **(B)** Outline of epimastigote transfection with Cas9/guide plasmid and linear homology template followed by limiting dilution at 24 hours post-transection (left). Schematic of the *TcCYP51* locus, homology-directed repair template, and predicted genomic locus following integration. The *TcCYP51* coding region and relevant homology arms are shown in orange, skip peptide in yellow, and the blasticidin-S deaminase gene in blue. Red triangles indicate the positions of stop codons. The purple horizontal line corresponds to the predicted binding location for the Southern blot probe; XhoI and XmaI sites in the genome are indicated. **(C)** Primers flanking *TcCYP51* were used to amplify the locus and identify clones with increased band size on an agarose gel. **(D)** Southern blot following digestion of genomic DNA with XhoI/XmaI and hybridization with a probe binding to the 3’ region of *TcCYP51*. **(E)** Construct design for genetic complementation with a C-terminal GFP tagged copy of *TcCYP51* and selection with puromycin. **(F)** Colocalization of GFP tagged CYP51 with the endoplasmic resident chaperone BiP. Mean Pearson’s correlation coefficient (PCC) between the BiP signal and the GFP signal of epimastigotes expressing CYP51::GFP is 0.55 (n = 707 parasites).

In other kinetoplastid parasites, the essentiality of CYP51 is species-specific; while *CYP51* deletion is tolerated in *Leishmania major* (Xu et al., 2014), deletion of *CYP51* in *Leishmania donovani* requires maintenance of an episomal copy of *CYP51* (McCall et al., 2015). Based mainly on the sensitivity of *T. cruzi* to azoles *in vitro* and results from early animal infection studies, it was concluded that CYP51 is essential for *T. cruzi* viability (Lepesheva et al., 2011).

In this study, we utilized a CRISPR/Cas9 strategy to disrupt *T. cruzi CYP51* translation and show that CYP51 activity is dispensable for the growth of *T. cruzi* epimastigotes. Loss of CYP51 activity leads to an accumulation of 14-methylated sterols with a concomitant absence of ergostane-type sterols. While *ΔCYP51* epimastigotes remain viable, their replication is slowed, and they exhibit extreme heat sensitivity and mitochondrial dysfunction. Disruption of the upstream enzyme, squalene epoxidase (SQLE) was sufficient to cause heat sensitivity and mitochondrial dysfunction suggesting that the loss of ergosterol and not the buildup of 14-methylated sterols in *ΔCYP51* epimastigotes is causative. Moreover, the inability of *ΔCYP51* epimastigotes to maintain the intracellular growth cycle in mammalian cells suggests the potential for contextual and stage-specific consequences of inhibiting *T. cruzi* sterol synthesis.

## Results

### Targeted gene disruption using Cas9 and homology mediated repair

To investigate the essentiality of CYP51, we designed a Cas9 mediated homology-directed repair strategy based on foundational work (Lander et al., 2015) to disrupt both *T. cruzi CYP51* alleles (TcCLB.510101.50; TcCLB.506297.260). Our modified approach utilizes Cas9 cut sites and homology-directed repair regions selected from conserved coding regions in *CYP51* (Figure 1B) to facilitate simultaneous integration at multiple allelic sites. The placement of an in-frame 2A skip peptide allows for translation of the selection marker followed by stop codons to prevent translation of the downstream *CYP51* sequence. This scheme creates a disruption of proper translation and consequently an ablation of function. Using a limiting dilution approach to clone parasites during the initial drug selection, we recovered parasites with insertions into both *CYP51* alleles, confirmed by PCR (Figure 1C) and Southern blot (Figure 1D). These parasites are referred to as *ΔCYP51* throughout. Genetic complementation was achieved using a modified pTREX expression vector encoding a C-terminally GFP tagged copy of *CYP51* (Figure 1E). Consistent with a role in sterol synthesis (Homma et al., 2000), CYP51-GFP localized predominantly to the endoplasmic reticulum as determined by colocalization with the ER chaperone BiP (Figure 1F).

### Sterol composition of ΔCYP51 epimastigotes

Using gas chromatography-mass spectrometry (GC-MS), we measured the effect of CYP51 disruption on free sterol composition in *T. cruzi* epimastigotes. An internal standard (ISTD) was used to compare the relative abundance of individual sterol species between samples. In line with previously published results, wild-type (WT) epimastigotes have detectable levels of exogenous (i.e., cholesterol) and endogenous sterols, including ergosterol (Figure 2). Similar to the effect of inhibiting CYP51 using ketoconazole, *ΔCYP51* epimastigotes completely lack sterols downstream of CYP51 and accumulate the 14-methylated sterol species, lanosterol, and eburicol. Genetic complementation led to the restoration of ergosterol synthesis and reduction in abundance, but not the elimination of eburicol (Figure 2). The lack of ergosterol in *ΔCYP51* parasites biochemically confirms the disruption of *CYP51* and demonstrates the dispensability of ergosterol for epimastigote proliferation and survival.

**Figure 2:**
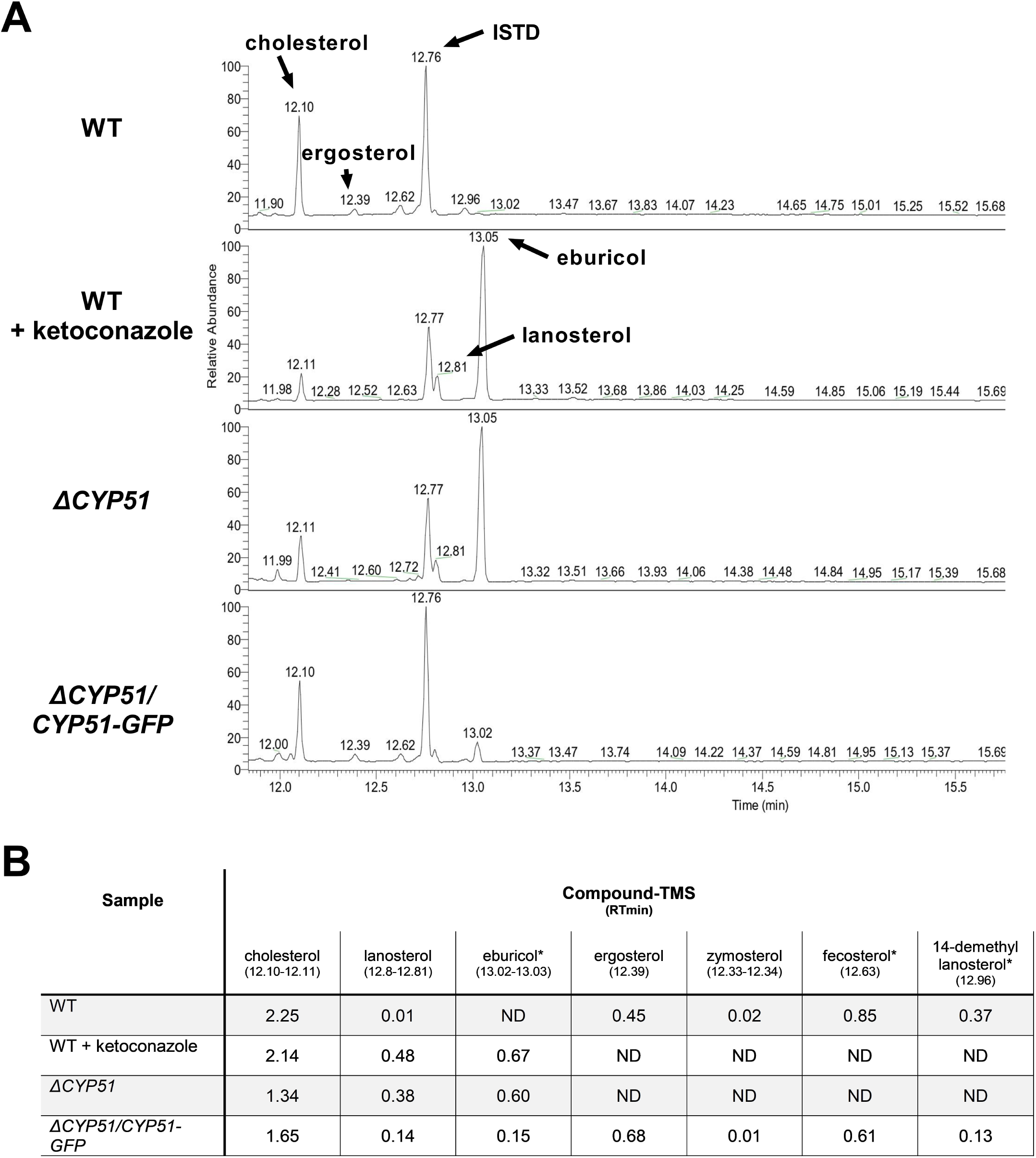
**(A)** GC-MS chromatogram of sterols derived from the indicated parasite lines. The internal standard (ISTD) along with endogenous and exogenously derived sterols are indicated. **(B)** Table of detectable sterol species and retention times from panel A and their relative signal compared to the ISTD. Sterol species were identified using a chemical standard unless indicated by an asterisk (*). ND = not detected. Ketoconazole was used at a final concentration of 4 μM.

### Quantification of parasite growth and sensitivity to anti-fungal compounds

The gene disruption and cloning strategy used allows for the isolation of parasites independent of their growth rate. Therefore, we quantified the doubling time of *ΔCYP51* parasites. In the exponential phase, WT epimastigotes double every 29.5 hours, and the doubling time of *ΔCYP51* parasites increases to 60.0 hours (Figure 3A). Genetic complementation partially restores the doubling time to 35.5 hours.

**Figure 3:**
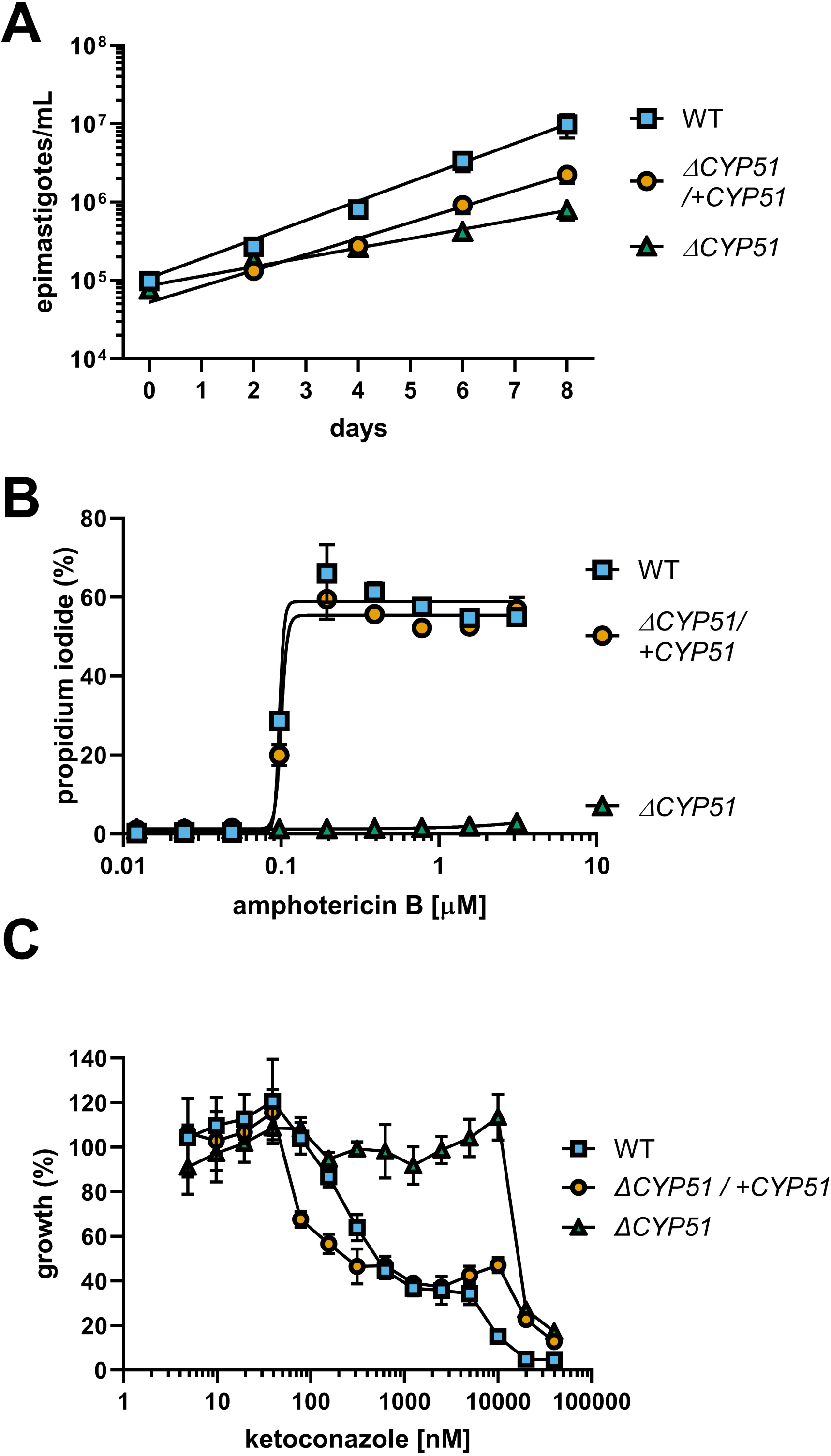
**(A)** Enumeration of epimastigote concentration every 48 hours starting at 1e5 epimastigotes/ml (n=3), mean ±SD shown. WT epimastigotes exited the logarithmic phase of growth after day 8. **(B)** Measurement of epimastigote membrane integrity by flow cytometry using propidium iodide staining following incubation for 4 hours with the indicated concentrations of amphotericin B (n=3), mean ±SD shown. **(C)** Growth of epimastigotes (8 days) relative to samples without ketoconazole treatment, starting at 1e5 epimastigotes/ml (n=3), mean ±SD shown.

Amphotericin B (ampB) activity is primarily dependent on binding to ergosterol and compromising the integrity of the plasma membrane (Gray et al., 2012). We measured the ability of ampB to compromise parasite membrane integrity through ergosterol binding. Consistent with the absence of ergosterol in the plasma membrane of *ΔCYP51* parasites, ampB does not compromise membrane integrity, unlike WT parasites, and sensitivity to ampB is restored in genetically complemented parasites (Figure 3B). Distinct from ampB, azoles such as ketoconazole inhibit an intermediate step in sterol synthesis mediated by CYP51. We found that at concentrations of ketoconazole up to ∼20 μM, *ΔCYP51* epimastigote growth is not inhibited, indicating a range where parasite growth inhibition by ketoconazole is solely dependent on CYP51 inhibition (Figure 3C). Conversely, at higher concentrations, *ΔCYP51* growth is inhibited, demonstrating secondary/off-target effects of ketoconazole at concentrations greater than 20 μM.

### Temperature sensitivity and mitochondrial dysfunction occur with CYP51 disruption

In addition to slowed growth and resistance to anti-fungal compounds, we hypothesized that similar to other pathogens (Hemmi et al., 1995; Xu et al., 2014; Mukherjee et al., 2020), alterations to sterol profiles may increase sensitivities of *T. cruzi* mutants to stress or metabolic dysfunction. Shifting the epimastigote growth temperature from 27 °C to 37 °C resulted in slowed growth of WT epimastigotes (Figure 4A). Unlike WT parasites, *ΔCYP51* epimastigotes fail to grow at 37 °C, and their decline in parasite numbers is accompanied by a precipitous rise in parasite death (Figure 4A).

**Figure 4:**
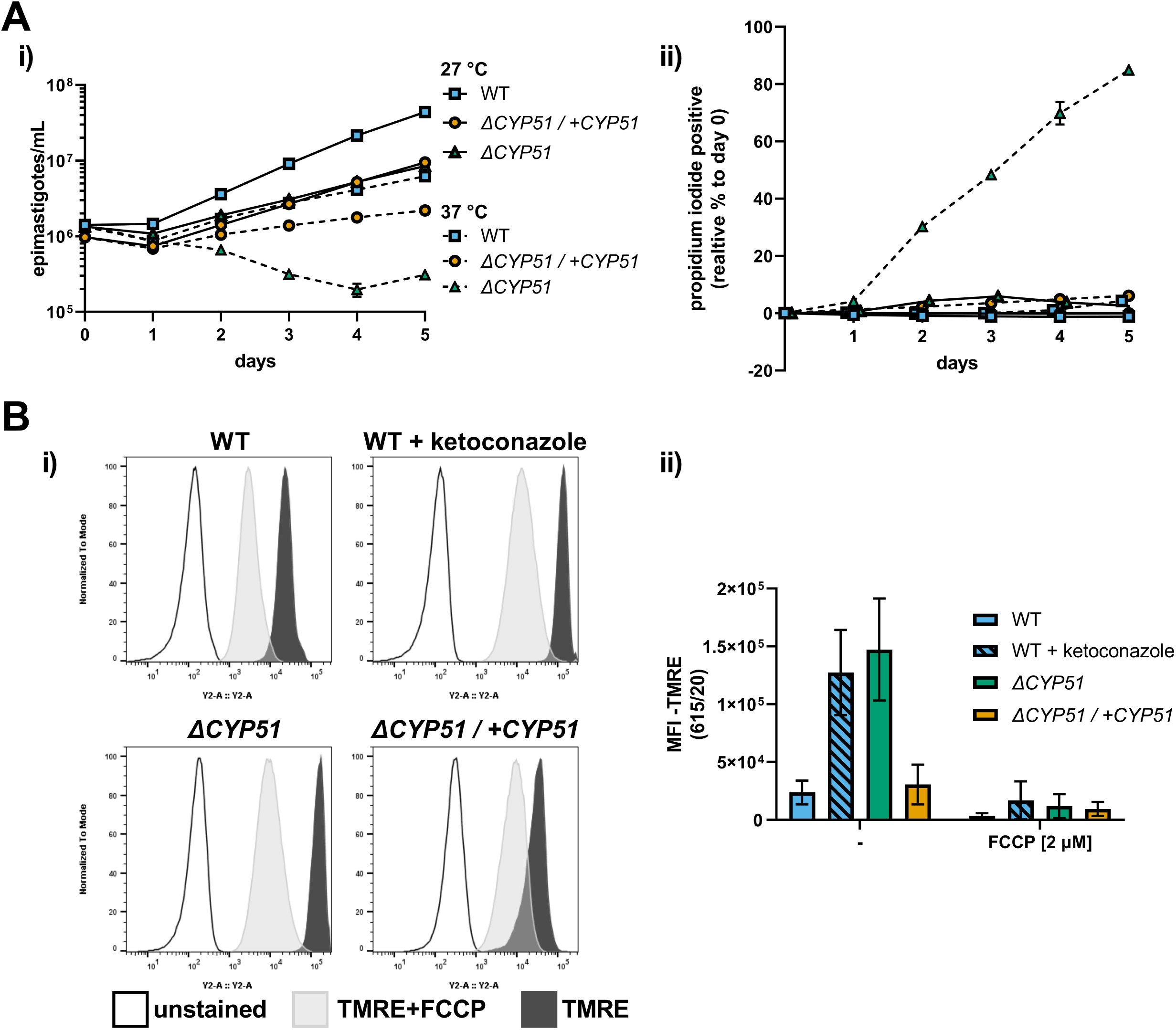
**(A)** (i) Growth and (ii) membrane integrity of epimastigotes assessed daily during continual exposure to either 27 °C or 37 °C (n=3), mean ±SD shown. **(B)** (i) Flow cytometry histograms of parasite fluorescence in the Y2 (615/20) channel. Unstained controls are shown along with TMRE stained parasites. (ii) Mean fluorescence intensity (MFI) in the Y2 channel with and without pre-treatment of parasites with FCCP, ±SD shown.

Sterols are integral membrane components in the plasma membrane and internal organelles, and alterations to sterol compositions can profoundly influence organelle function (Rodrigues et al., 2001; Mukherjee et al., 2020). We measured the mitochondrial membrane potential using tetramethylrhodamine ethyl ester (TMRE), a dye that accumulates in active mitochondria proportional to the mitochondrial membrane potential. We discovered that *ΔCYP51* parasites have an increased mitochondrial membrane potential (Figure 4B) and that under nutritional stress, mitochondrial oxidative stress is increased (Supplementary Figure 1).

### Conversion of ΔCYP51 epimastigotes to the mammalian stages in vitro is impaired

Genetic manipulation of *T. cruzi* is currently limited to the epimastigote stage. Consequently, to study the impact of any genetic modification on the mammalian stages of *T. cruzi*, it is necessary to generate host cell invasive metacyclic trypomastigotes from epimastigotes, which is routinely achieved *in vitro*. The commonly used method of inducing metacyclogenesis in triatomine artificial urine (TAU) medium resulted in the death of *ΔCYP51* epimastigotes (data not shown). However, we find that metacyclic trypomastigotes were formed in both WT and *ΔCYP51* stationary epimastigote cultures at varying efficiencies (Supplementary Figure 2A) and could be purified using a charge-based separation method (Cruz-Saavedra et al., 2017). However, *ΔCYP51* metacyclic trypomastigotes were poorly infective (Supplementary Figure 2B), and the intracellular amastigotes observed at 18 hours post-infection failed to persist and replicate *in vitro* under the conditions tested (Supplementary Figure 2C).

### Disruption of squalene epoxidase is sufficient to cause temperature sensitivity and mitochondrial dysfunction

Loss of CYP51 activity leads to the simultaneous buildup of 14-methylated sterols and absence of ergosterol. To distinguish between effects of the accumulation of 14-methylated sterols and loss of ergostane type sterols we targeted squalene epoxidase (SQLE, TcCLB.509589.20; TcCLB.503999.10), the enzyme that catalyzes the conversion of squalene to 2,3-oxidosqualene. SQLE activity is essential for sterol synthesis; in other systems disruption of SQLE leads to the accumulation of squalene and sterol auxotrophy (Garcia-Bermudez et al., 2019). We targeted *SQLE* using a CRISPR/Cas9 in both WT and *ΔCYP51* parasites (Figure 5A). In both backgrounds, we obtained clones with integration into all *SQLE* alleles, confirmed by PCR (Figure 5B) and Southern blot (Figure 5C).

**Figure 5:**
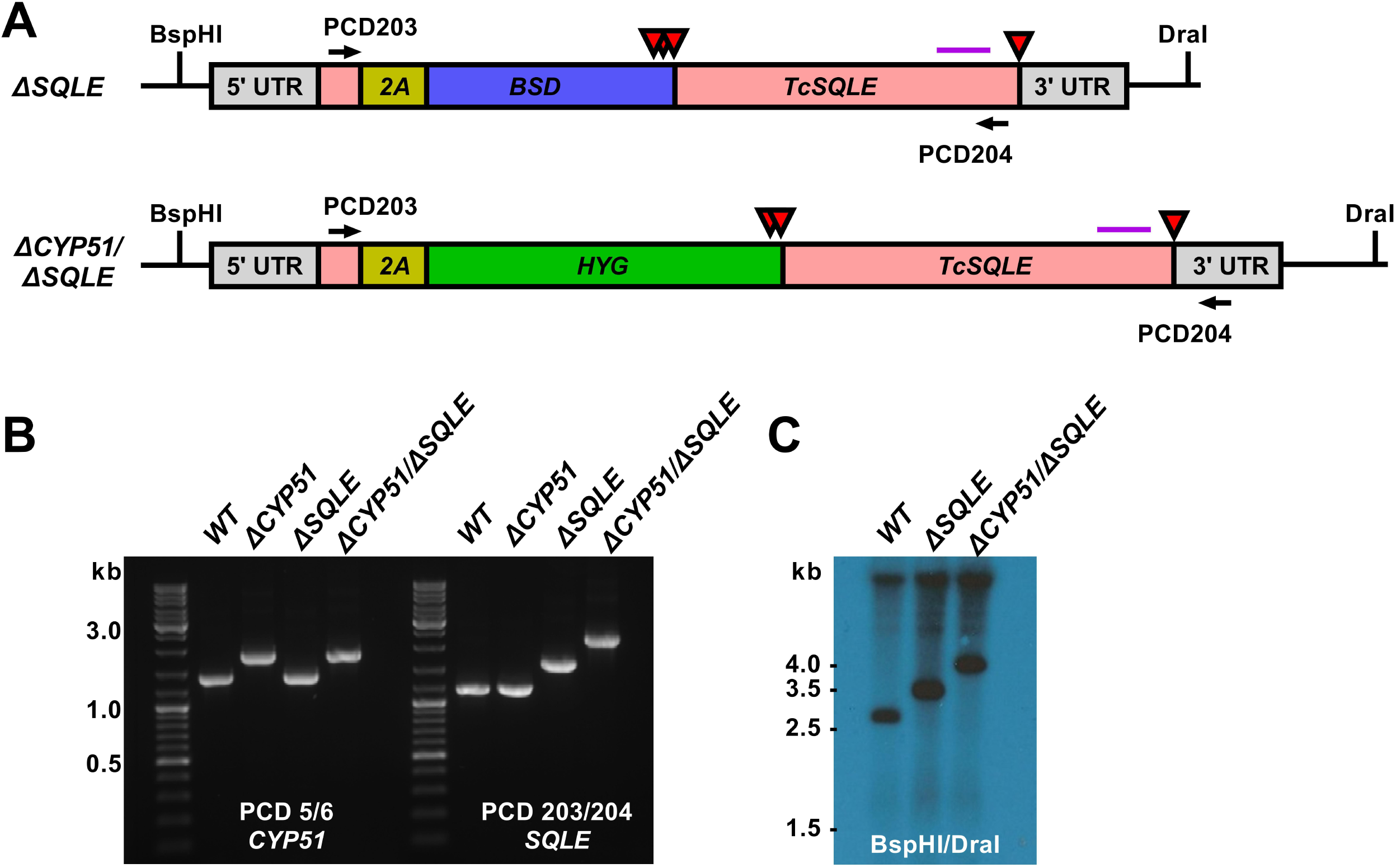
**(A)** Predicted *TcSQLE* loci following homology-directed repair template integration shown. The *TcSQLE* coding region is shown in pink, skip peptide in yellow. Blasticidin-S deaminase (blue) was used in WT parasites while hygromycin B phosphotransferase (green) was used for transfection of *ΔCYP51* parasites already carrying the blasticidin-S deaminase gene. Red triangles indicate the positions of stop codons. The purple horizontal line corresponds to the predicted binding location for the Southern blot probe; BspHI and DraI sites in the genome are indicated. **(B)** Primers flanking *TcSQLE* were used to amplify the locus and identify clones with increased band size on an agarose gel. **(D)** Southern blot following digestion of genomic DNA with BspHI/DraI and hybridization with a probe binding to the 3’ region of *TcSQLE*.

Consistent with an essential role of SQLE in endogenous sterol synthesis we found that *ΔSQLE* and *ΔCYP51/ΔSQLE* parasites lack endogenous sterols, including 14-methylated species (Figure 6). Both *ΔSQLE* and *ΔCYP51/ΔSQLE* parasites have a concomitant increase in squalene, the substrate of SQLE (Figure 6). The only detectable sterol species in *ΔSQLE* and *ΔCYP51/ΔSQLE* parasites is exogenous cholesterol (Figure 6B). Similar to *ΔCYP51* parasites the loss of SQLE and absence of ergosterol leads to absolute resistance to ampB (Figure 7A).

**Figure 6:**
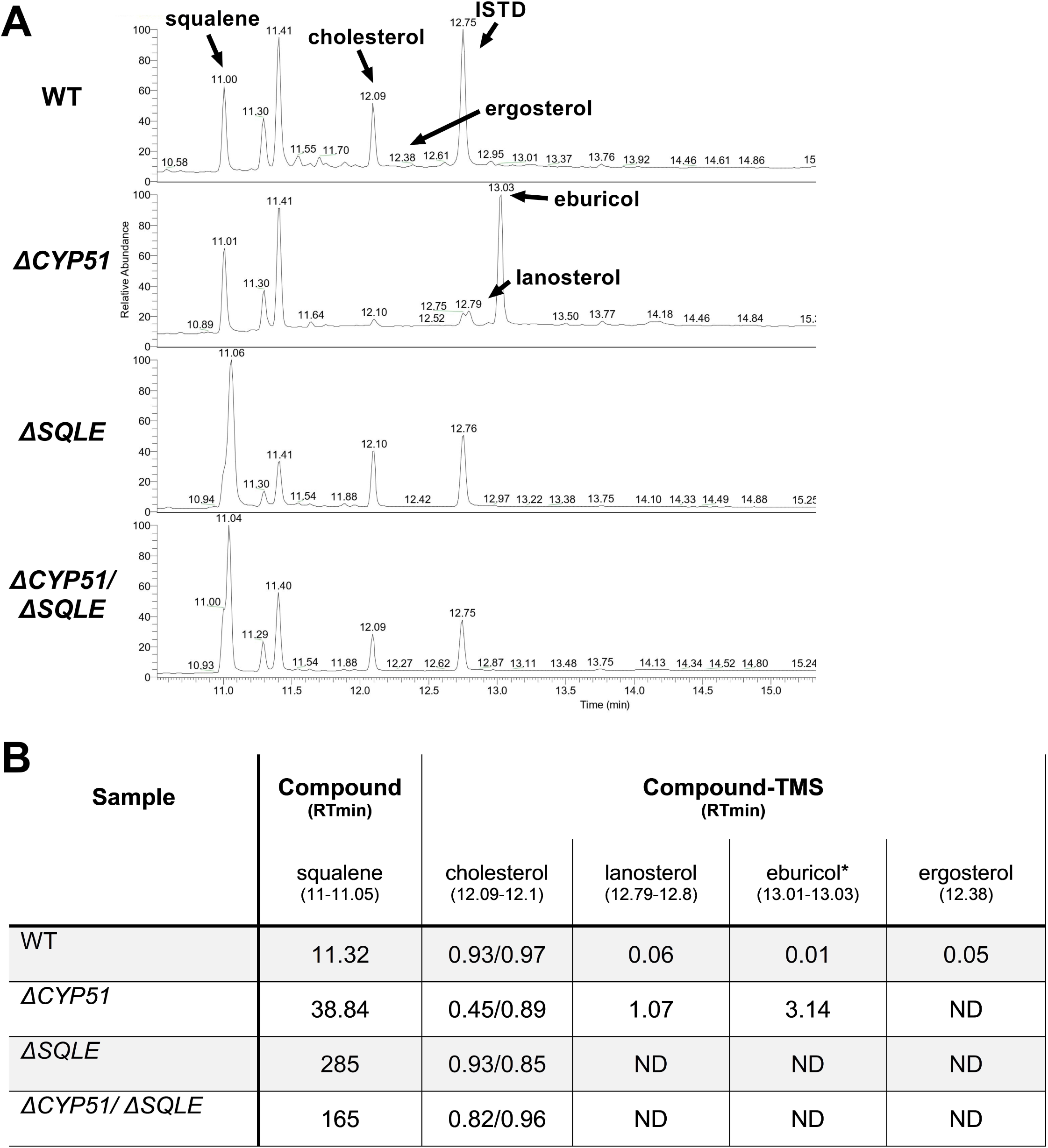
**(A)** GC-MS chromatogram derived from lipid extractions from the indicated parasite lines. The internal standard (ISTD) along with endogenous and exogenously derived sterols are indicated along with squalene. **(B)** Table of detectable sterol species and retention times from panel A and their relative signal compared to the ISTD. Each sample was split and half underwent saponification (2 hours at 68 °C in 1M methanolic potassium hydroxide) to cleave cholesterol esters. ISTD normalized values are shown for cholesterol with and without saponification (free sterols/post saponification). Species were identified using a chemical standard unless indicated by an asterisk (*). ND = not detected.

**Figure 7:**
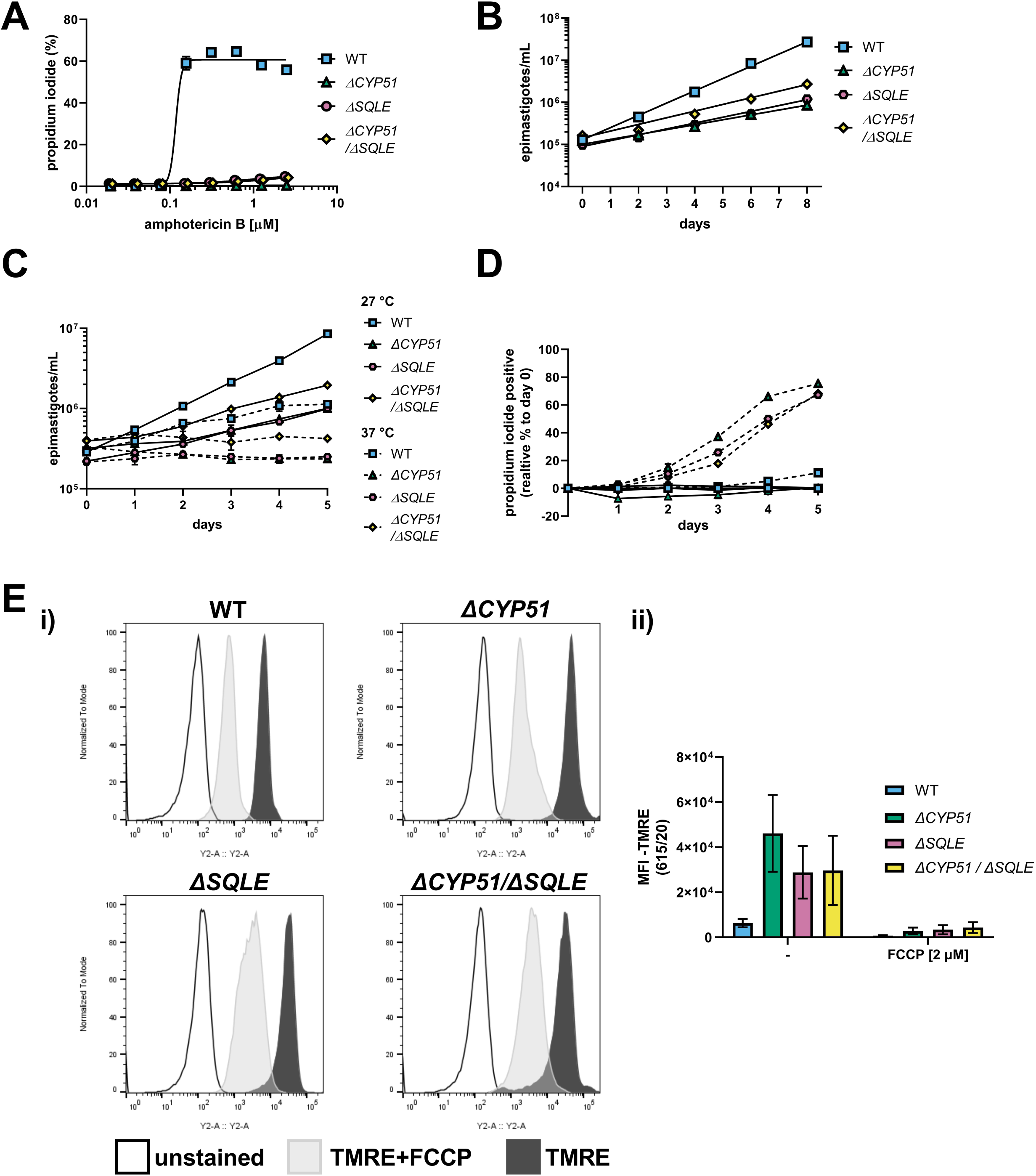
**(A)** Measurement of epimastigote membrane integrity by flow cytometry using propidium iodide staining following incubation for 4 hours with the indicated concentrations of amphotericin B (n=3), mean ±SD shown. **(B)** Measurement of epimastigote concentration every 48 hours starting at 1e5 epimastigotes/ml (n=3), mean ±SD shown. **(C)** Growth and **(D)** membrane integrity of epimastigotes assessed daily during continuous exposure to either 27 °C or 37 °C (n=3), mean ±SD shown. **(E)** (i) Flow cytometry histograms of parasite fluorescence in the Y2 (615/20) channel. Unstained controls are shown along with TMRE stained parasites. (ii) Mean fluorescence intensity (MFI) in the Y2 channel with and without pre-treatment of parasites with FCCP ±SD shown.

Both *ΔSQLE* and *ΔCYP51/ΔSQLE* epimastigotes have an increased doubling time relative to WT; 52.0 and 45.7 hours respectively vs. 24.8 hours (Figure 7B). However, *ΔCYP51* doubling time remained the most protracted at 62 hours (Figure 7B). Similar to *ΔCYP51* epimastigotes, SQLE deficient parasites failed to proliferate at 37 °C (Figure 7C) and lost their membrane integrity over time (Figure 7D). *ΔSQLE* parasites were also incapable of maintaining mitochondrial membrane potential at levels similar to WT parasites (Figure 7E) suggesting that cholesterol is insufficient to maintain optimal organelle membrane functions and oxidative stress (Supplementary Figure 3).

## Discussion

Similar to fungi, the major products of sterol biosynthesis in *Trypanosoma cruzi* and other trypanosomatid species are ergostane-type sterols which require sterol 14α-demethylase (CYP51) for their production (Lepesheva et al., 2011). The sensitivity of *T. cruzi* to azoles, potent inhibitors of CYP51 used to treat fungal infections, led to the assumption that ergosterol synthesis is essential for *T. cruzi* growth and survival (Lepesheva et al., 2018; Osorio-Méndez and Cevallos, 2018). Here, we provide the first demonstration that *CYP51* can be genetically disrupted in *T. cruzi* epimastigotes without loss of parasite viability. Consistent with the effects of pharmacological inhibition of CYP51 on the sterol composition of *T. cruzi* (Ottilie et al., 2017; Dumoulin et al., 2020), the *ΔCYP51* mutant lacks ergosterol and accumulates 14-methylated sterols; lanosterol and eburicol. While there are no gross morphological defects associated with these changes in sterol composition, *ΔCYP51* epimastigotes proliferate more slowly than WT and exhibit heightened sensitivity to elevated temperature, resulting in parasite death. Moreover, the marked increase in mitochondrial membrane potential and elevated mitochondrial superoxide levels observed in the *ΔCYP51* mutant is consistent with the recognized impact of altered sterol composition on mitochondrial physiology and function (Rodrigues et al., 2001; Mukherjee et al., 2020). Thus, while CYP51 is not essential for *T. cruzi* epimastigote growth, our data suggest that the sterol 14α-demethylase activity of this enzyme remains important for mitigating mitochondrial stress and for resilience to thermal shifts in this parasite.

Notably, the *T. cruzi ΔCYP51* mutant is incapable of establishing and maintaining the mammalian infection cycle *in vitro* under the conditions tested. The transition from the epimastigote stage to mammalian cell-invasive metacyclic trypomastigotes and subsequently to intracellular replicative amastigotes involves significant morphological and metabolic remodeling and exposure to elevated temperatures and low pH. Given the reduced tolerance of the *ΔCYP51* mutant to thermal stress, it is not surprising that key developmental transitions needed to establish mammalian cell infection are compromised in the mutant. Yet, recent studies show that azole-treated *T. cruzi* amastigotes can proliferate under certain growth conditions despite the lack of ergostane-type sterols and the accumulation of 14-methylated sterols (Dumoulin et al., 2020). Azoles also do not provide sterile cure *in vivo* (Molina et al., 2014; Torrico et al., 2018). These findings demonstrate that the requirement for CYP51 can be context-dependent, with important implications for the outcome of therapeutic treatment. Therefore, it is possible that conditions may exist that allow for growth of the *ΔCYP51* mammalian stages.

In other kinetoplastid protozoan parasites, the requirement for CYP51 expression can be species-specific. For example, CYP51 is essential for *Leishmania donovani* (McCall et al., 2015) but dispensable for *Leishmania major* (Xu et al., 2014; Mukherjee et al., 2020). Strikingly, many of the phenotypic characteristics described for *T. cruzi ΔCYP51* epimastigotes are similar to those reported for CYP51-deficient *L. major* promastigotes, which also exhibit growth defects and sensitivity to environmental stressors (Xu et al., 2014; Mukherjee et al., 2020). In *Leishmania*, the impairment of CYP51 does not result in an accumulation of lanosterol or eburicol; rather, sterol biosynthesis proceeds without removing the C14 methyl group, resulting in 14-methyl fecosterol and 14-methyl zymosterol (Xu et al., 2014). Since these C14 species are more similar to ergosterol, it has been suggested that they may more readily substitute for ergosterol in membranes (Lepesheva et al., 2018), thereby allowing these parasites to exist without CYP51 activity. However, we find that *T. cruzi* epimastigotes can proliferate without further modification of lanosterol and eburicol.

Sterols are relatively minor components of phospholipid bilayers, but their presence dramatically alters the biophysical properties of membranes, including fluidity and permeability (Cournia et al., 2007). The loss of ergosterol and the accumulation of 14-methylated sterols in *ΔCYP51* parasites occurs in tandem, and therefore separating the individual impact of these outcomes is challenging. Characterization of a *L. major* sterol biosynthesis mutant lacking sterol methyltransferase (SMT) activity revealed the expected loss of ergostane-type sterols but without the production of 14-methylated sterols due to the presence of functional CYP51 (Mukherjee et al., 2019, 2020). Unlike CYP51-deficient parasites, *L. major ΔSMT* promastigotes do not display heightened temperature sensitivity (Mukherjee et al., 2019, 2020), highlighting the potential link between 14-methylated sterol accumulation and temperature sensitivity in these parasites. To address the role of 14-methylated sterols in *T. cruzi* stress tolerance we targeted squalene epoxidase. SQLE is essential for sterol synthesis but acts upstream of sterol ring formation at the conversion of squalene to 2,3-oxidosqualene. Loss of SQLE activity leads to the accumulation of squalene and absence of endogenous sterols leading to sterol auxotrophy. We found that both WT and *ΔCYP51* parasites remained viable following disruption of SQLE. *ΔSQLE* parasites lacked all endogenous sterols including any 14-methylated species, suggesting that exogenous cholesterol is sufficient to fulfill the parasite’s sterol requirement; however, *ΔSQLE* parasites grow more slowly than WT parasites and have decreased tolerance to elevated temperature along with mitochondrial dysfunction. The similarity of these phenotypes with *ΔCYP51* parasites implies that the lack of ergosterol and not the buildup of 14-methylated sterols most prominently influences epimastigote stress tolerance.

The maintenance of sterol homeostasis has been extensively studied in mammals and yeast, where intricate feedback mechanisms regulate sterol uptake and synthesis (Goldstein et al., 2006). Little is known about the regulation of sterol pools in kinetoplastid protozoan parasites, but studies in *T. brucei* have shown that lipoprotein (LDL-cholesterol) uptake is associated with reduced sterol synthesis, suggesting that feedback systems may exist in these parasites as well (Coppens and Courtoy, 1995; Rodrigues et al., 2001). Both the extracellular and intracellular stages of *T. cruzi* incorporate exogenous cholesterol into membranes. We find that relative to WT *T. cruzi* epimastigotes, *ΔCYP51* parasites display a relative reduction in free cholesterol content, potentially due to parasite regulation of total free sterol content, including the buildup of intermediates. Consistent with the influence of 14-methylated sterols on free cholesterol we found that cholesterol levels returned to WT amounts in *ΔCYP51/ΔSQLE* epimastigotes. Additionally, saponification of *ΔCYP51* lipid extracts led to an increase in free cholesterol demonstrating that levels of free sterols can be regulated through the maintenance of cholesterol esters.

Taken together, these data suggest that endogenous sterol synthesis is conditionally essential in *T. cruzi* epimastigotes. Similar observations have been made regarding the conditional essentiality of CYP51 activity in amastigotes (Dumoulin et al., 2020). The ability of epimastigotes to fulfill their sterol requirement entirely through scavenging is unexpected and further demonstrates that metabolic flexibility can contribute to parasite resilience in diverse environments (Dumoulin and Burleigh, 2018, 2021). Since the presence of endogenous sterols is important for *T. cruzi* stress tolerance, it is likely that this pathway has been maintained to allow for parasite resilience in diverse environments but is dispensable under certain growth conditions. The sensitivity of *T. cruzi* mammalian stages to azoles *in vivo*, without the ability to provide sterilizing cure, may reflect this conditional importance of endogenous sterol synthesis (Molina et al., 2014; Khare et al., 2015; Torrico et al., 2018).

Additionally, partner drugs or therapies should be explored that take advantage of altered parasite biology caused by inhibition of sterol synthesis. In *Leishmania major* CYP51 knockout parasites, the increased mitochondrial membrane potential leads to the hypersensitization of parasites to pentamidine, likely due to elevated accumulation in the mitochondrion (Mukherjee et al., 2020). Therefore, continued studies characterizing *T. cruzi* sterol biology will be instrumental in understanding the design of therapies aimed at eliminating parasites.

## Materials and Methods

### Parasite Culture

*Trypanosoma cruzi* Tulahuén LacZ clone C4 was obtained from the American Type Culture Collection (ATCC, PRA-330) and propagated as epimastigotes at 28 °C in filter-sterilized liver infusion tryptose (LIT) consisting of 4 g/L NaCl, 0.4 g/L KCl, 8 g/L Na2HPO4, 2 g/L dextrose, 3 g/L liver infusion broth, 5 g/L tryptose, with 25 mg/L hemin and supplemented with 10% heat-inactivated fetal bovine serum (FBS). LIT was supplemented with a chemically defined lipid mixture (Sigma, St. Louis, Missouri) containing non-animal derived fatty acids and cholesterol during epimastigote cloning.

### Cas9 Gene Editing and Selection

#### Single plasmid expression of Cas9 and sgRNA

The plasmids pTREX-n-Cas9 and pUC_sgRNA (Lander et al., 2015) were gifts from Dr. Roberto Docampo (Addgene plasmid 68708 and 68710). We first modified pTREX-n-Cas9 by removing the HA and GFP tags while maintaining two copies of the SV40 NLS at the C-terminus of Cas9 (Supplementary Table 1). Guide RNAs were identified using the Eukaryotic Pathogen gRNA Design Tool (EuPaGDT) (Peng and Tarleton, 2015) and selected based on their predicted absence of secondary targets and ability to target multiple alleles. A sgRNA sequence to target *GFP* was amplified by PCR using pUC_sgRNA as a template and cloned into pTREX-n-Cas9 using the BamHI restriction sites as described (Lander et al., 2015). Targeting *GFP* (pTREX-n-Cas9-gGFP3) was used to optimize transfection and selection (data not shown). Altering sgRNA specificity was performed by site-directed mutagenesis and pTREX-n-Cas9-gGFP3 as a template (Supplementary Table 1).

#### Generation of template for homology-directed repair and gene disruption

The template for generating homology-directed repair DNA for gene disruption was constructed by inserting a P2A viral skip peptide in frame with a downstream blasticidin-S deaminase (BSD) or hygromycin B phosphotransferase gene (HYG) containing three or two stop codons respectively and TOPO cloned into a pCR4 backbone (Thermo Fisher, Waltham, Massachusetts). Linear homology-directed repair donor DNA was generated by PCR using ultramers (IDT, Coralville, Iowa) containing 100-bp of homology to regions flanking the predicted Cas9 cut site.

#### Transfection of pTREX-n-Cas9 and HDR template

For transfection, pTREX-n-Cas9-sgRNA (CYP51_94 and CYP51_99rc or SQLE_117 and SQLE_111rc) and homology template were precipitated and re-suspended in a small volume of water (<15 µl / transfection). Epimastigotes (∼4e7) were washed and re-suspended in 100 µl/transfection of a nucleofection solution consisting of 90 mM NaPO_4_, 5 mM KCL, 0.15 mM CaCl_2_ and 50 mM HEPES at pH 7.3 (Schumann Burkard et al., 2011). Epimastigotes in nucleofection solution were mixed with DNA, transferred to a disposable 2 mm gap cuvette (BTX Harvard Apparatus, Holliston, MA), and transfected using an Amaxa Biosystems Nucleofector II (Lonza, Cologne, Germany) set to program U-33. Transfected parasites were re-suspended in 10 ml LIT and allowed to recover for 24 hours prior to cloning by limiting dilution in 96-well plates in the presence of blasticidin (10 µg/ml) or hygromycin (350 µg/ml). Between 40-60 days post-transfection, wells were visually inspected for parasites and selected for growth in upright T-25 flasks, and screened for cassette insertion by PCR (Supplementary Table 1).

### Southern Blot

Genomic DNA was extracted from epimastigotes and digested overnight with the indicated restriction enzymes. Digested DNA (5 µg/well) was separated on a 1% agarose gel. The gel was then covered in depurination solution (HCl 250 mM) until bromophenol blue in the sample loading buffer changed from blue to yellow, approximately 60 minutes. After removing the depurination solution, the gel was washed three times with distilled water and covered with a denaturation solution (NaCl 1.5M, NaOH 0.5M) for 60 minutes, followed by three washed with distilled water. The gel was then submerged in a neutralization solution (NaCl 1.5M, Tris HCl 0.5M, pH 7.5) for 50 minutes. DNA was transferred to a Hybond-N+ membrane (GE Amersham, Amersham, UK) overnight by capillary blot. Following an overnight transfer, the membrane was dried and UV crosslinked using a GS Gene Linker UV chamber (Bio-Rad, Hercules, CA). Probes derived from PCR products (Supplementary Table 1) were labeled using the Amersham ECL™ Direct Nucleic Acid Labeling and Detection System (GE Amersham, Amersham, UK) and hybridized as described in the manufacturer’s instructions, followed by exposure to film.

### Flow Cytometry

Epimastigotes were analyzed using a MACS Quant VYB (Miltenyi Biotec, Bergisch Gladbach, Germany) equipped with a 405 nm, 488 nm, and 561 nm laser. Mitochondrial membrane potential was measured using a tetramethylrhodamine ethyl ester (TMRE) and carbonyl cyanide 4-(trifluoromethoxy) phenylhydrazone (FCCP) as a control (Abcam, Cambridge, UK). Prior to incubation and acquisition, epimastigotes were diluted to 4e6 parasites/ml and stained with a final concentration of 100 nM TMRE and, where indicated, pre-exposed to 5 µM FCCP. Quantification of mitochondrial superoxide production was performed using MitoSOX™ Red (Thermo Fisher, Waltham, Massachusetts). Excitation of TMRE and MitoSOX™ Red utilized the yellow laser (561nm) and emission in the Y2 channel (615/20). DAPI (Thermo Scientific, Waltham, MA (1:100 dilution, 1 mg/ml stock) was used exclude dead parasites.

For the determination of parasite counts (e.g., growth measurements), epimastigotes were collected and fixed in 1% PFA or run live when measured with propidium iodide. Sample concentrations were diluted when necessary to fall within the linear range of the instrument (<10,000 events per second). Stop gates were not used; the entire uptake volume was acquired to reflect total parasite concentrations in a given uptake volume.

### Sterol Extraction and Identification by GC-MS

Sterols were extracted and identified as described previously (Sharma et al., 2017; Dumoulin et al., 2020). Prior to sterol extraction, sitosterol-d7 (Avanti Polar Lipids, Alabaster, Alabama) was added as an internal standard (ISTD) at 1.12 μg/2e7 epimastigotes. Epimastigote cell pellets were extracted three times with C:M (2:1, v/v) and centrifuged each time at 1800 x g for 15 minutes at 4 °C. The supernatant was dried under a constant stream of N_2,_ and the resulting material was subjected to a Folch’s partitioning (4:2:1.5, C:M:W). Following centrifugation, the lower phase from the Folch’s partitioning was dried under N_2_ and re-suspended in chloroform before being passed over a silica 60 column. BSTFA + 10% TMCS/pyridine (5:1, v/v) was added to each sample, vortexed, and heated at 70 °C for 30 minutes prior to injection.

### Immunofluorescence microscopy

*Trypanosoma cruzi* epimastigotes expressing CYP51::GFP or Δ*CYP51* epimastigotes were fixed in 1% paraformaldehyde and permeabilized with TritonX-100. Parasites were placed on poly-L-lysine coated slides and stained with a rabbit α-BiP primary antibody (gift from Jay Bangs, 1:1000 dilution) (Bangs et al., 1993), and a goat α-rabbit secondary antibody conjugated to Alexa Fluor 647 (Invitrogen, Waltham, MA, 1:1000 dilution). DAPI (Thermo Scientific, Waltham, MA (1:5000 dilution, 1 mg/ml stock) was used to identify parasite DNA. Parasites were mounted in ProLong Diamond (Thermo Fisher, Waltham, Massachusetts) and cured for 24 hours. Parasites were imaged on a Yokogawa CSU-X1 spinning disk confocal system paired with a Nikon Ti-E inverted microscope and an iXon Ultra 888 EMCCD camera. The 100x lens was used for imaging, and image processing, analysis, and display were completed in FIJI (Schindelin et al., 2012).

### Colocalization analysis

Colocalization metrics were analyzed using the ImageJ plugin EZColocalization (Stauffer et al., 2018); all default settings were used unless otherwise noted here. The channels corresponding to BiP (known ER protein) and CYP51::GFP were input into EzColocalization, using a duplicated BiP channel as a mask for cell identification. The Pearson’s correlation coefficient (PCC) was calculated for all parasites in the images. As a control for the method, a PCC was also calculated for the Δ*CYP51* parasite line (PPC = 0.55).

## Figure Legends

**Supplementary Figure 1: (A)** Mean fluorescence intensity (MFI), ±SD, measured by flow cytometry of live parasites stained with mitSOX red mitochondrial superoxide for 1 and **(B)** 6 hours. Staining and incubation solutions are indicated.

**Supplementary Figure 2: (A)** Percentage of metacyclic trypomastigotes as assessed by Giemsa staining following a five-day incubation in DMEM supplemented with 2% FBS. **(B)** Invasion, persistence, and **(C)** replication of metacyclic trypomastigotes following DEAE sephacel enrichment and infection of dermal fibroblasts at a multiplicity of infection of 5 parasites/host cell. Host cells were fixed with PFA and stained with DAPI prior to counting the number of amastigotes per cell at the indicated time points; median indicated in red. ND = none detected.

**Supplementary Figure 3:** Mean fluorescence intensity (MFI), ±SD, measured by flow cytometry of live parasites stained with mitSOX red mitochondrial superoxide for 6 hours.

## Conflict of Interest

The authors declare that the research was conducted in the absence of any commercial or financial relationships that could be construed as a potential conflict of interest.

## Author Contributions

PCD - Conceptualization, Experimentation, Data analysis, Supervision, Funding acquisition, Writing

JV - Experimentation, Data analysis

MW - Experimentation, Data analysis

JXW – Experimentation (GC-MS)

BAB - Conceptualization, Data analysis, Supervision, Funding acquisition, Writing

## Funding

The study was supported in part by NIH NIAID R21AI146815 grant awarded to BAB and AHA 19POST34380209 awarded to PCD.

## Acknowledgments

The α-BiP antibody was a gift from Jay Bangs, The State University of New York at Buffalo.

## Notes

### Competing Interest Statement

The authors have declared no competing interest.

