## Supplementary figures and images for "Endogenous sterol synthesis is dispensable for *Trypanosoma cruzi* epimastigote growth but not stress tolerance"

### Supplementary Figure 1

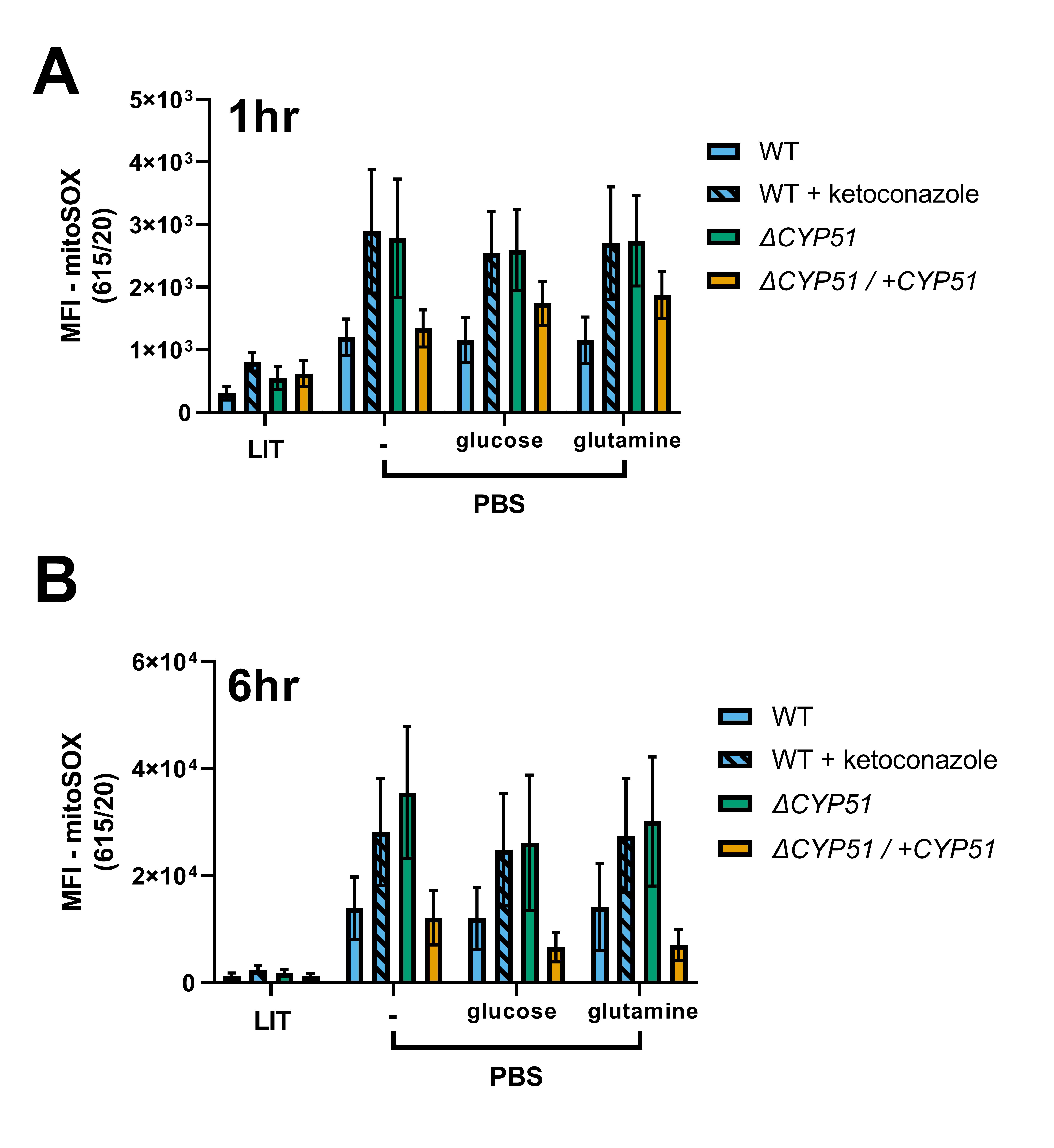

### Supplementary Figure 2

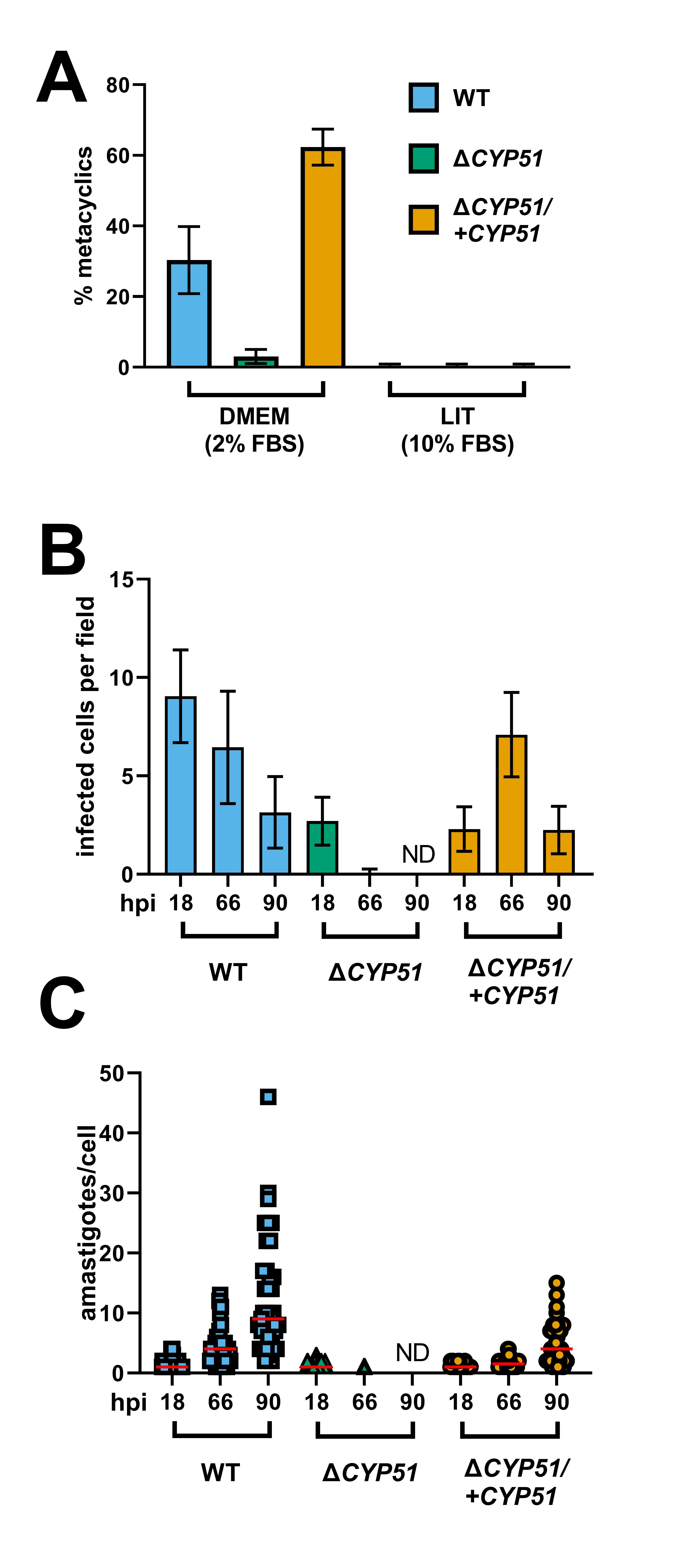

### Supplementary Figure 3

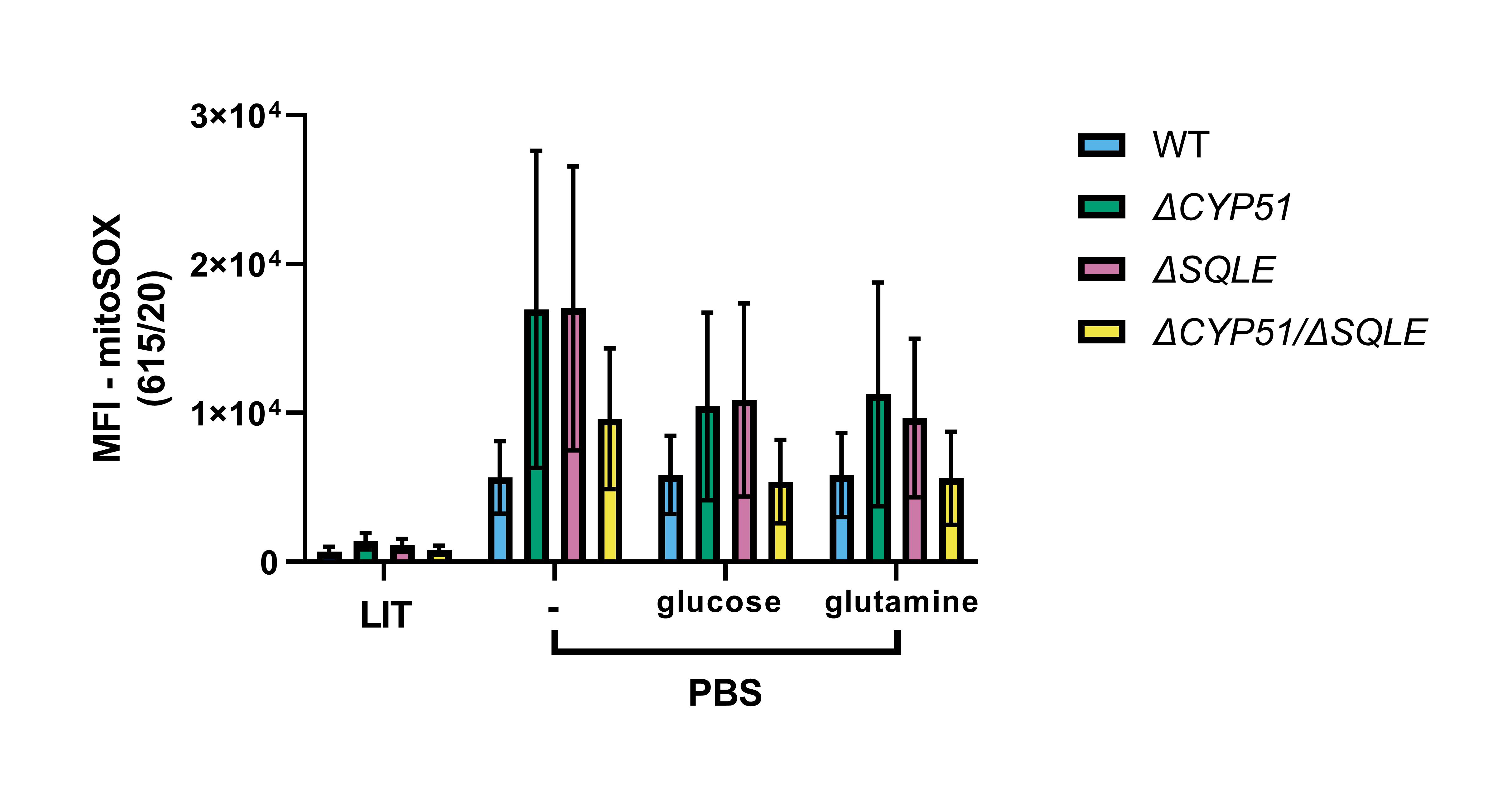
