## Supplementary Table 1 for "Endogenous sterol synthesis is dispensable for *Trypanosoma cruzi* epimastigote growth but not stress tolerance"

| Name | 5' – 3' sequence | Note |
|---|---|---|
| Primers for modification of pTREX-Cas9 (HA tag removal) | | |
| Q5SDM_F | CCCAAAAAGAAAAGGAAGGTTGATTAGAAGCTTATCGATACCGTCGAC | |
| Q5SDM_R | GTCCTCGACTTTTCGCTTCTTTTTCGGGTCGCCTCCCAGCTGAGA | |
| sgRNA targeting specificity | | |
| CYP51_99rc | TCCCAGAAACGGCACCGTAA | |
| CYP51_94 | CTGGGACACATTGTGCAGTT | |
| SQLE_117 | AGGTGGAAGCATCGCTGGAC | |
| SQLE_111rc | GATGGCATCGTAATCGTAAT | |
| Primers for generation of universal homology directed repair (HDR) template | | |
| PCD111 | GCTACTAACTTCAGCCTGCTGAAGCAGGCTGGCGACGTGGAGGAGAACCCT GGACCTATGGCCAAGCCTTTGTCTCAAGA | P2A-BSD HDR template_F |
| PCD112 | TGGCGGCCGCTCTAGAACTAGTGGATTTAGCCCTCCCACACATAAC | P2A-BSD HDR template_R |
| PCD92 | GCTACTAACTTCAGCCTGCTGAAGCAGGCTGGCGACGTGGAGGAGAACCCT GGACCTATGAAAAAGCCTGAACTCAC | P2A-HYG HDR template_F |
| PCD93 | TGGCGGCCGCTCTAGAACTAGTGGAT | P2A-HYG HDR template_R |
| Ultramer primers for generation of HDR DNA by PCR | | |
| PCD119 | ATTGAAGCCATTGTATTGGCCCTTACGGCTCTCATCCTGTACTCGGTGTACTC TGTAAAGTCATTTAACACAACCCGTCCTACTGACCCACCGGTTTACGCTACTA ACTTCAGCCTGCT | CYP51_F |
| PCD120 | ATCGCCTACGATCGTCACGCGCTGTCCCCCAATGCTGATGGTGAAAACACCC GACTTGAGTTCACGCTTGCATCGCTGCATAAACTCAAGCGGGTTCTTGCTAG GCGGCCGCTCTAGAACTAGTGGA | CYP51_R |
| PCD209 | ATGTTCTGCACTTTCACTTTGCTGGTGGTGGTCACCGTACTGATACTCAATCA CGTACTCTCTCGACTGCGGTTTAAGCCCACCCGCGCTACTAACTTCAGCCTGC T | SQLE_F |
| PCD210 | GGCTGCAAGAGTTCACCGACAATGCGATCCGGTTTGACAAAGAGAGAACGC TCAAGCAGTAGAACCTTCCGATTTTGTTCTGAAAGCGCTTTTGCCATCCTAGG CGGCCGCTCTAGAACTAGTGGA | SQLE_R |
| Southern blot probe generation | | |
| PCD7 | GCGATCCCCACGAACACAGTCG | CYP51_F |
| PCD8 | TTGTTCTTTGGATGCATCAGAT | CYP51_R |
| PCD211 | ATTATGTGTGACGGCGGCTCCT | SQLE_F |
| PCD212 | CAAGACTGCGGGGCAATGACGT | SQLE_R |
| Primers for genotyping epimastigote clones | | |
| PCD5 | ACGGCTCTCATCCTGTACTCGG | CYP51_F |
| PCD6 | CGGGTGTACTTCACAAGACACT | CYP51_R |
| PCD203 | ATGTTCTGCACTTTCACTTTGC | SQLE_F |
| PCD204 | ACGTCATTGCCCCGCAGTCTTG | SQLE_R |
